# NCPepFold: Accurate Prediction of Non-canonical Cyclic Peptide Structures via Cyclization Optimization with Multigranular Representation

**DOI:** 10.1101/2024.12.05.626948

**Authors:** Qingyi Mao, Tianfeng Shang, Wen Xu, Silong Zhai, Chengyun Zhang, An Su, Chengxi Li, Hongliang Duan

## Abstract

Artificial intelligence-based peptide structure prediction methods have revolutionized biomolecular science. However, restricting predictions to peptides composed solely of 20 natural amino acids significantly limits their practical application, as such peptides often demonstrate poor stability under physiological conditions. Here, we present NCPepFold, a computational approach that can utilize a specific cyclic position matrix to directly predict the structure of cyclic peptides with non-canonical amino acids. By integrating multi-granularity information at the residue- and atomic-level, along with fine-tuning techniques, NCPepFold significantly improves prediction accuracy, with the average peptide RMSD for cyclic peptides being 1.640 Å. In summary, this is a novel deep learning model designed specifically for cyclic peptides with non-canonical amino acids without length restrictions, offering great potential for peptide drug design and advancing biomedical research.

## 1. Introduction

In recent years, peptides have emerged as a promising class of potential therapeutics.^[1,2]^ Compared to small-molecule drugs, peptides possess distinct physicochemical properties, including larger sizes and more flexible backbones, which offer inherent advantages as inhibitors or activators of protein-protein interactions (PPIs).^[3–5]^ Cyclic peptides, in particular, exhibit superior stability and bioactivity compared to linear peptides.^[6–8]^ Given the significant potential of peptides in the therapeutic market,^[9]^ understanding their structure and internal interaction is crucial to advance novel peptide design. The rise of deep learning (DL) has led to remarkable progress in biomolecular modeling.^[10–13]^ AlphaFold2 (AF2) and RoseTTAFold have achieved unprecedented accuracy in protein structure prediction,^[14,15]^ delivering accuracy that rivals the resolution of crystallography experiments. Research demonstrates that, despite AF2’s training set excluding peptides shorter than 16 residues, it successfully learned key protein folding mechanisms from millions of data points. This allows AF2 to make remarkably accurate predictions of peptide-protein complexes, as peptide binding can be seen as a natural extension of protein folding.^[16]^ Building on this understanding, AfCycDesign introduced cyclic constraints into the AF2 framework to embed circular information for cyclic peptide structure prediction.^[17]^ HighFold further improved accuracy by integrating structural information on head-to-tail circle and disulfide bridge, enhancing the accuracy of disulfide bond prediction while ensuring proper cyclization.^[18]^

Despite these encouraging findings, there remain limitations in practical drug development when reconciling idealistic approaches with real-world challenges. A key limitation of current models is their focus on peptides composed solely of natural amino acids. While such peptides have therapeutic potential, their real-world applications remain limited due to low stability against proteases and poor bioavailability.^[19]^ However, by introducing non-natural amino acids, the pharmacokinetic properties of peptides can be significantly improved.^[20]^ One notable example is Selepressin, a cyclic peptide composed of nine amino acids, in which the fourth and eighth positions are substituted with two non-canonical amino acids.^[21,22]^ This vasopressin derivative maintains similar target selectivity while offering a significantly extended plasma half-life, making it a more effective therapeutic option. Moreover, the weak forces in peptides, such as hydrogen bonds, van der Waals forces, and hydrophobic interactions, are often insufficient to stabilize secondary structure conformations. Introducing non-canonical amino acids with side-chain modifications can mimic the structures of natural products or hotspots in PPIs, thereby stabilizing the peptide’s secondary structure. However, progress in structure prediction for peptides with non-canonical amino acids has been slow thus far. This is primarily due to the wide diversity of non-canonical amino acids, making it impractical to treat them in the same manner as the 20 standard amino acids. The absence of a suitable representation method and an accompanying network framework remains a significant challenge. Additionally, the limited number of high-resolution structures for these peptides makes it challenging to train a new peptide-specific DL model from scratch. Alternatively, new DL models could be trained on a large set of peptide structures with non-canonical amino acids generated by Rosetta.^[23]^ However, the accuracy and performance of a model would be inherently limited by the quality of the data used to generate the training set. ADCP is a peptide-protein docking method capable of docking linear peptides or cyclic peptides with non-canonical amino acids.^[24]^ However, it is only effective for peptides with fewer than 30 amino acids and requires the structural information of the target protein, which sometimes does not meet our needs.

Recently, RoseTTAFold All-Atom (RFAA), an all-atom version of RoseTTAFold, has been introduced.^[25]^ Building upon RoseTTAFold’s network framework, RFAA effectively incorporates information related to chemical modifications. By modeling modified residues and chemical groups as atom-bond graphs, it combines sequence-based description with atomic graph representation. In protein glycosylation structure predictions, RFAA achieved a median modification RMSD of 3.2 Å. Modeling non-canonical amino acids is similar to that of covalent modifications. However, we speculate that RFAA’s training set likely excluded peptide samples with fewer than ten residues, as the model shows low prediction accuracy for peptides with around ten residues. Moreover, among the millions of training data in RFAA, there are only 4,775 protein-multi-residue ligand complexes (including protein-glycan and protein-peptide). Due to data scarcity and the different data distributions between peptides and proteins, the model, while proficient in learning protein structural information, lacks a comprehensive understanding of peptide structures. Furthermore, RFAA is unable to effectively handle cyclic peptides, which restricts its scope of application.

Here, we present NCPepFold, a deep learning model specifically designed for peptides with non-canonical amino acids. By extending the feature construction module of RFAA, NCPepFold is capable of accurately predicting the structures of peptides containing non-canonical amino acids (including monomers and complexes) based solely on sequence information (**Figure 1a**). Specifically, we integrated a Transformer-based peptide sequence model, leveraging bidirectional self-attention to atomize residues and capture interactions between residues and atoms.^[26]^ Structural information is embedded within the coevolutionary data of residues. By incorporating multilevel representations in the structural feature module, NCPepFold effectively captures the inherent hierarchical relationships within peptides, leading to a more sophisticated and rational peptide learning framework. This integrated approach significantly enhances the model’s ability to analyze the multi-level details of amino acids, particularly non-canonical amino acids. For cyclic peptides, we incorporated a cyclization module, introducing Cyclic Position Matrix (CPM) into the relative position matrix of residues. In addition, we fine-tuned NCPepFold to enable transfer learning from protein folding mechanisms to peptide structural biology. In conclusion, our NCPepFold achieves high-precision modeling of both linear and cyclic peptides with non-canonical amino acids, as well as monomers and complexes.

**Figure 1.**
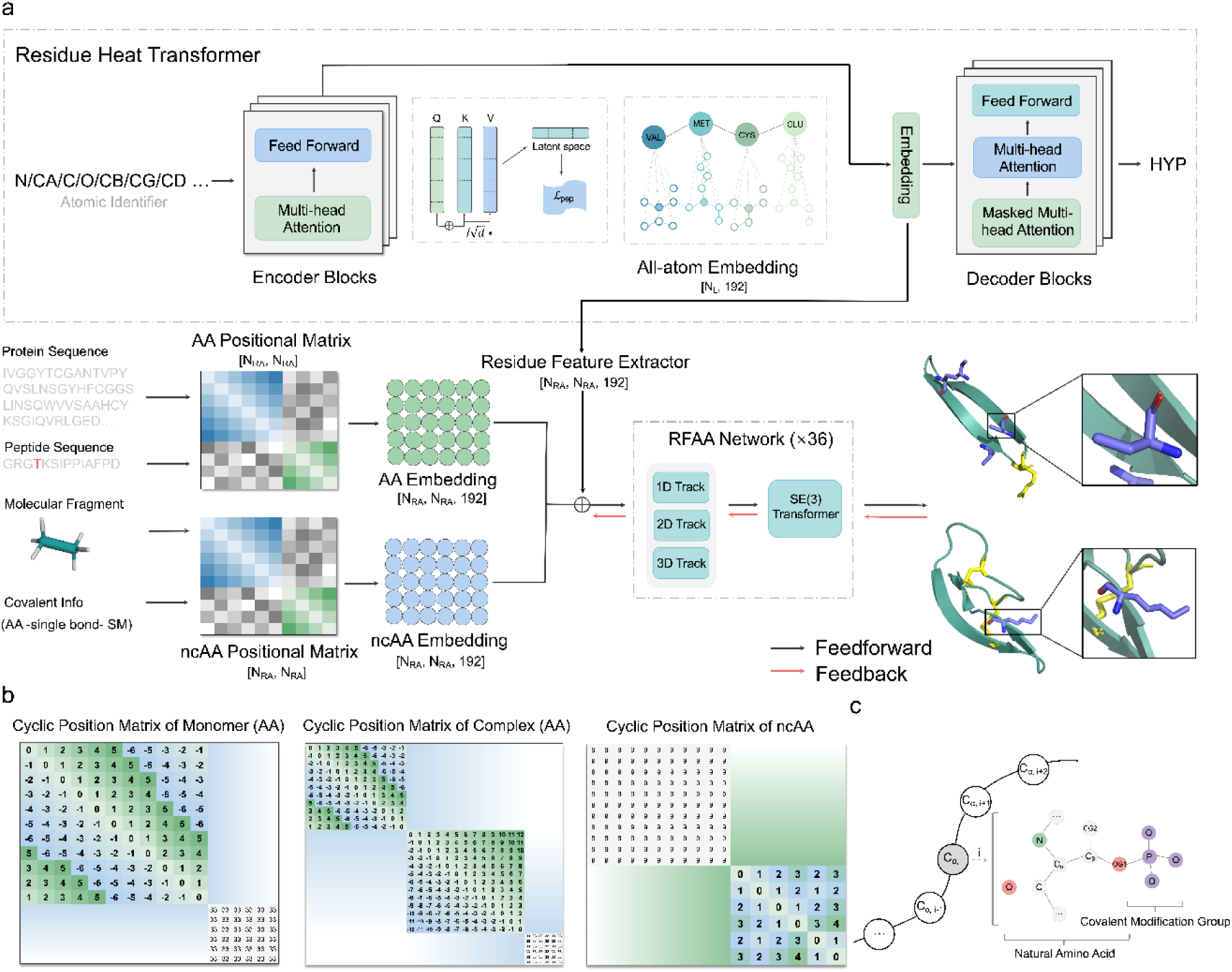
The overview of the NCPepFold to predict peptide structure with non-canonical amino acids. a) Method overview. The network consists of a feature construction module and a prediction module. The feature construction module introduces Residue Heat Transformer (RHT) block, a Transformer-based model. The output of RHT encoder, which has dimensions [1, N_L_, 192] where N_L_ is the total number of atoms in the peptide chain, passes through the Residue Feature Extractor to align its dimensions with those of two relative position encoding feature embeddings. And for cyclic peptides, a cyclic constraint is added to the relative position matrix. On the right is the visualization of predicted structures, with purple indicating non-canonical amino acids and yellow highlighting the cyclic regions. b) CPM, tailored for head-to-tail cyclic peptides. The left shows the relative position matrix of standard amino acids in cyclic peptide monomers with non-canonical amino acids; the middle displays the same matrix for cyclic peptide complexes; and the right presents the atomic adjacency matrix for the non-canonical amino acids in both monomers and complexes. c) represents non-canonical amino acid representation. NCPepFold splits non-canonical amino acid into standard amino acid and side-chain molecular groups, which are connected by covalent bonds.

In this work, we evaluated NCPepFold ‘s accuracy in predicting the structures of cyclic and linear peptides with non-canonical amino acids with secondary structures available in RCSB Protein Data Bank (PDB). We evaluated the performance of NCPepFold by calculating two types of RMSD (peptide RMSD and modification RMSD), DockQ, and the fraction of native contacts (F_nat_). The specific calculation methods for various evaluation metrics can be found in Methodology.

## 2. NCPepFold Methodology

### 2.1. NCPepFold Framework

Figure 1a illustrates the architecture of NCPepFold and the information flow during model training and inference. The model input consists of three components: amino acid sequences of the protein and peptide, molecular graph of the modification group, and the insertion position of the modification. NCPepFold represents peptides containing non-canonical amino acids as a combination of sequence description and atomic bond graph. Non-canonical amino acids are extracted from the sequence and atomized together with the modification group, generating a relative position matrix for the standard amino acid residues and an atomic relative position matrix for the non-canonical amino acids. For cyclic peptides, CPM algorithm is applied to the standard amino acid relative position matrix derived from linear peptides, with cyclization information further incorporated. After incorporating the all-atom information from Residue Heat Transformer module into two relative position matrices mentioned above, these inputs are passed into SE3-Transformer to predict atomic coordinates.

### 2.2. Network Deployment

#### 2.2.1. Cyclic Position Matrix

We modified the relative position matrix of peptides for the structural prediction of cyclic peptides containing non-canonical amino acids. CPM encodes both the forward (N-to-C) and backward (C-to-N) relationships between each residue in the cyclic peptide. For peptide monomers, CPM is applied to the entire input sequence, while for protein-peptide complexes, CPM is only applied to the peptide portion, with the protein sequence retaining the default matrix. This design ensures model can distinguish structural differences between peptides and proteins. For cyclic peptides containing non-canonical amino acids, residue relative position matrix’s size becomes (N-x)-(N-x), where N is residue number and x is non-canonical amino acid number. Although some amino acids are excluded, the sequence length in CPM formula remains the same, ensuring matrix accurately reflects cyclic conformation of the peptide.

CPM is expressed as:

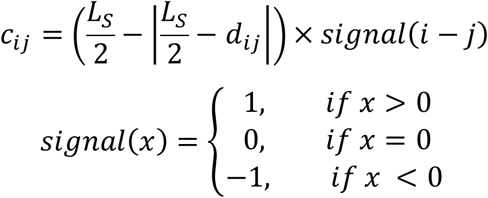

where *c*_*ij*_ denotes the relative position between residue *i* and residue *j* in the relative position cyclic matrix, *d*_*ij*_ denotes the relative position between residue *i* and residue *j* in the linear matrix, *L*_*S*_ denotes the number of residues (including the non-canonical residues), |·| denotes the symbol of absolute value. The sign function, Signal(x), is used to represent the relative spatial relationships between residues in a cyclic peptide sequence. Specifically, it indicates relative direction of amino acids. *Signal* = 1 means amino acid *i* is located to the right of amino acid *j*, *Signal* = −1 means amino acid *i* is located to the left of amino acid *j*, and *Signal* = 0 means both amino acids are at the same position. This function helps capture the topological features of the cyclic peptide (Figure 1b), providing spatial representations for NCPepFold.

#### 2.2.2. Residue Heat Transformer

In feature construction module, we incorporated atomic interaction information of residues to enhance the mapping relationship between residues and their corresponding atoms. Specifically, we introduce a Residue Heat Transformer (RHT) (Figure 1a), an attention-based neural network architecture, which uses multi-head attention layers to learn different levels of peptide information. RHT encoder consists of multiple layers of multi-head self-attention (MHA) and feedforward networks. After integrating positional information, the feature data is fed into a six-layer RHT encoder module, providing a global perspective of the entire peptide chain. To enhance the generalization capability of RHT, the dropout trick is applied within the MLP. The output embeddings of RHT are passed through an MLP to predict potential residue distributions, and the generated feature representations are then projected into the target dictionary using an activation function, mapping the real-valued vectors accordingly. Both the encoder and decoder modules incorporate 8 heads Multi-Head Self-Attention blocks, focusing on the relationships between the residues. The relevant formula is as follows:

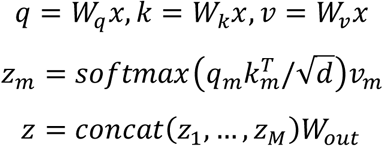

NCPepFold ‘s initial stage involves pre-training RTH to provide a well-initialized state for joint training, aiding faster convergence. This pre-training is performed in a self-supervised manner using a unique masked language modeling objective. To accommodate variable-length batch inputs, each residue embedding is padded to a fixed shape with zeros. Subsequently, the RTH module, containing prior knowledge, is merged with the main module for end-to-end joint training, where atomic representations are aggregated at the amino acid level, mixed with amino acid features extracted by the main module, and together inputted into the atomic coordinate prediction module.

#### 2.2.3. Fusion of RFAA and RHT

To train RHT in NCPepFold, we devised a specialized algorithm that aligns the learned atomic interaction information with the positional feature information of standard and non-canonical amino acids. For the RHT, we extract the encoder output 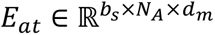, where *b*_*s*_ is the batch size during model training, *N*_*A*_ is the total number of atoms in the target peptide chain, and *d*_*M*_ is the embedding dimension of the network. This represents the attention features generated by the Transformer, which encapsulate high-dimensional information about the input atomic features, reflecting the interactions and positional relationships between atoms. To enable to integrate with the embedding of the relative position matrix, we reshape *E*_*at*_ using a Mapping block. Specifically, for standard amino acids, we aggregate atomic-level representations to amino acid-level representations 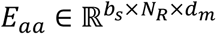, where *N*_*R*_ is the number of standard amino acid. For non-canonical amino acids, the representation retains its original shape. For these two types, we compute pairwise features between residues and atoms using Einstein summation. Finally, through Block Diagonal Concatenation, we combine amino acid pairwise features and atomic pairwise features, resulting in an 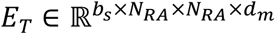 that matches the dimensionality of the embedding of NCPepFold. Here, *N*_*RA*_ is the total number of standard amino acids and the total number of atoms in non-canonical amino acids, and this is ultimately merged with the embedding of the relative position matrix.

#### 2.2.4. Predicted Coordinates Processing

In the predicted three-dimensional coordinates *xyz*_*pred*_ of NCPepFold, the atomic coordinates of non-canonical amino acids are positioned at the end. To align the structure during the loss computation phase, we rearrange and pad the predicted atomic coordinates to ensure that the non-canonical amino acids are inserted in the correct positions and match the expected structural format. Specifically, we adjust *xyz*_*pred*_ according to the atomic position indices *I* from the ground truth, moving non-canonical amino acids from the end to their correct locations. Furthermore, we identify the maximum number of atoms *N*_*max*_ across all amino acid sequences and create a padding array based on this value to ensure dimensional consistency in the output tensor. This reordering function guarantees that the atomic coordinates correspond accurately to their respective amino acids or structural units in subsequent analyses and visualizations.

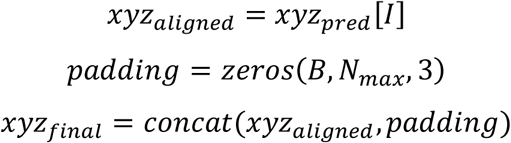

## 3. Experimental Section

### 3.1. Datasets

#### 3.1.1. Training datasets

We constructed a diverse training dataset, which consists of three subsets: a head-to-tail cyclic peptide dataset, a dataset of peptides cyclized by disulfide bridges, and a linear peptides dataset which contains non-canonical amino acids. Each subset is further divided into monomers and complexes. The cyclic peptide dataset was obtained from HighFold, containing 63 cyclic peptide monomers with experimental structures and 34 cyclic peptide complexes.^[18]^ The disulfide-bonded cyclic peptide monomer dataset contains 46 samples, sourced from CPPsite,^[27]^ while the complex dataset contains 62 samples from Propedia.^[28]^ The linear peptide dataset containing non-canonical amino acids was extracted from the test set provided by Singh’s work on their website, which, after curation, resulted in 32 monomers and 174 complexes as part of the training set.^[29]^ This dataset is implemented to enable the model to comprehensively learn the features of different types of peptide structures. More details about datasets can be seen in Section S1 of Supporting Information.

#### 3.1.2. Test datasets

In test set, we focus on peptides with secondary structures, as these structures confer stability to peptides, reducing the likelihood of amide bond hydrolysis by enzymes in the body and enhancing the drug-likeness of peptides.

The structures in the test set were all downloaded from the PDB database. Except for the head-to-tail cyclic peptides, the remaining samples have a secondary structure content greater than 50%, calculated using DSSP.^[30]^ This was because the data for head-to-tail cyclic peptides with non-canonical amino acids are scarce, and strict filtering would result in an insufficient sample size for statistical analysis. Test set includes 18 linear peptide monomers, 22 linear peptide complexes, 10 head-to-tail cyclic peptide monomers, and 23 disulfide-cyclized peptide monomers.

During the preprocessing of PDB files, for files containing multiple states, we retained the first state. For the convenience of processing, all solvents, water molecules, and heteroatoms were removed from the structures. For atoms with isomeric states, we retained the coordinates of state A. In the case of homodimers, one set of chains was extracted for subsequent analysis. In our work, non-canonical amino acids were split into standard amino acids and modification groups when constructing profiles. In complexes, we defined the shortest chain as peptide chain and labeled it as chain A for ease of subsequent analysis.

### 3.2. Loss Function

NCPepFold requires two stages of training: pre-training of RTH, aimed at providing a rational initialization for joint training and accelerating the convergence process; and fine-tuning of NCPepFold. For RTH pre-training, loss function includes a cross-entropy term to maximize learning of atomic interaction information. For end-to-end joint training, we use frame aligned point error (FAPE) for the main chain as the loss function, which better preserves the overall structural topological consistency.^[31]^

Cross-entropy loss function is as follows:

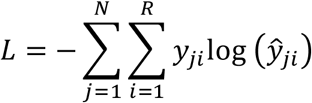

FAPE loss function is as follows:

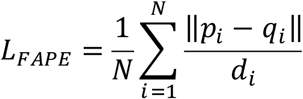

where *p* and *q* represent the coordinate sets of the main chain atoms in the predicted and true structures, respectively, while *d* is a normalization factor used to adjust the relative importance of the error.

### 3.3. Evaluation metrics

When evaluating model performance, certain metrics are required as benchmarks. Here, we used peptide RMSD (or ligand RMSD for complex) and modification RMSD to assess the accuracy of predicted structures, while DockQ and F_nat_ were employed to evaluate the interactions between peptide-protein complex.

#### 3.3.1. RMSD

RMSD measures the average deviation in distance between corresponding atoms in the native and predicted structures. In this work, we used peptide RMSD to represent the overall deviation of the peptide chain and modification RMSD to assess the accuracy of the local structure of non-canonical amino acids.

To evaluate the overall deviation of the peptide chain, we calculated peptide RMSD. For predicted structures, we skipped residues and atoms that were missing in the native structure, and for the native structure, we ignored atoms that are missing in the predicted structure (such as OXT). Next, we extracted the *C*_*⍺*_ atoms from both the native and predicted structures, aligning the coordinates using Kabsch alignment algorithm.^[32,33]^ For monomers, we aligned all *C*_*⍺*_ atoms in the peptide chain and calculated *C*_*⍺*_ RMSD of the entire chain. For peptide-protein complex, peptide RMSD was defined similarly to ligand RMSD, where, after aligning the protein’s *C*_*⍺*_ atoms, *C*_*⍺*_ RMSD of the peptide chain was calculated.

To assess the deviation between the local structure of non-canonical amino acids and their corresponding native structures, we calculated modification RMSD. The modification RMSD, also referred to as RMSD of non-canonical amino acid, is defined similarly to the modification RMSD in RFAA. First, the atoms within 10 Å of the non-canonical amino acid are aligned, and then calculate all-atom RMSD for the non-canonical amino acid. If a peptide chain contains multiple non-canonical amino acids, we first calculate the modification RMSD for each individually, then compute the overall average.

Specifically, the RMSD calculation is expressed as follows:

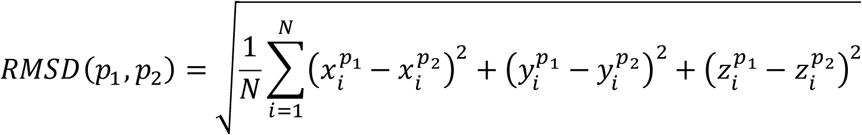

where *p*_1_ denotes peptide conformation from chemical experiments, *p*_2_ denotes peptide conformation predicted by NCPepFold, N denotes length of peptide chain and index i traverses the number of *C*_*⍺*_ atoms. The variables *x*, *y* and *z* correspond to the 3D coordinates of each residue.

#### 3.3.2. DockQ

To evaluate the docking quality of peptide-protein complexes, we used DockQ to assess the overall quality of the predicted structures.^[34]^ DockQ is a score ranging from 0 to 1, combining three metrics: F_nat_, LRMS, and iRMS. Based on the DockQ score, the structural quality of the complex is classified into four categories: ‘Incorrect’ (0.00 ≤ DockQ < 0.23), ‘Acceptable’ (0.23 ≤ DockQ < 0.49), ‘Medium Quality’ (0.49 ≤ DockQ < 0.80), and ‘High Quality’ (DockQ ≥ 0.80).

DockQ calculation is expressed as follows:

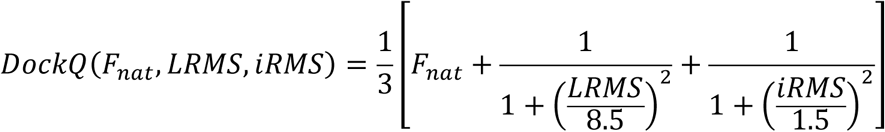

#### 3.3.3. Fnat

When dealing with peptide-protein complexes, peptides tend to bind to shallow grooves on the protein surface, resulting in many atoms that do not interact directly with the protein. This can lead to inflated RMSD, even though the peptide-protein interaction itself may be accurate. To more precisely capture these interactions, as shown in other peptide-related studies,^[24,35]^ we introduced F_nat_ to represent docking success rate.^[36]^ Native contacts are defined as peptide-protein residue pairs where the heavy atoms are within 5 Å in the crystal complex. F_nat_ ranges from 0 to 1, with peptide-protein complexes having F_nat_ ≥ 0.5 considered to be of high docking quality.

## 4. Results

### 4.1. Accurate prediction of cyclic peptide structure

We first evaluated NCPepFold on two test sets: 10 cyclic peptide monomers with modified residues forming head-to-tail cyclization from the PDB and 23 cyclic peptides with disulfide bonds. As shown in **Table 2** and **Table 3**, NCPepFold demonstrated strong predictive performance, with both the mean and median values for peptide RMSD and modification RMSD < 2 Å, which is considered indicative of high-quality predictions.

**Table 1.** Composition of test set.

| Name | Length | NCAA number | Secondary structure |
| --- | --- | --- | --- |
| Linear peptide monomer | 10-25 | 1-2 | $\alpha$ helix |
| Linear peptide complex | 12-36 | 1-3 | $\alpha$ helix |
| Head-to-tail cyclic peptide monomer | 10-34 | 1-6 | $\beta$ sheet |
| Disulfide-bonded cyclic peptide monomer | 12-41 | 1-3 | $\alpha$ helix, $\beta$ sheet, mixed structure |

**Table 2.**
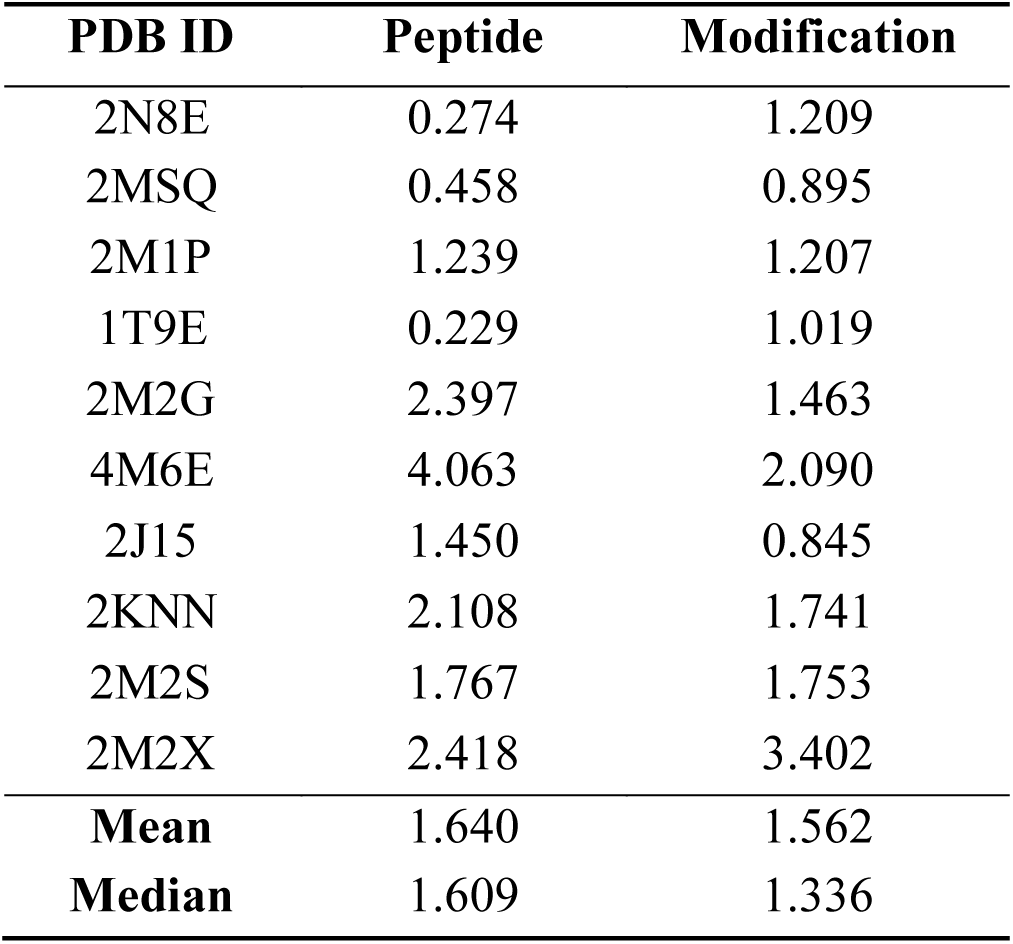
RMSD results of head-to-tail cyclic peptides.

**Table 3.** RMSD results of disulfide-cyclized peptides.

| **PDB ID** | **Peptide** | **Modification** | **PDB** | **Peptide** | **Modification** |
| --- | --- | --- | --- | --- | --- |
| 1P9G | 2.104 | 2.301 | 2MFX | 1.824 | 3.118 |
| 2CRD | 1.203 | 1.313 | 2MG6 | 1.637 | 3.283 |
| 1BIG | 1.252 | 1.490 | 3L09 | 0.474 | 1.218 |
| 1K64 | 1.740 | 1.132 | 4E86 | 0.763 | 0.907 |
| 2EW4 | 3.587 | 1.558 | 4E83 | 1.075 | 0.963 |
| 6MY3 | 1.684 | 1.484 | 3HJD | 0.632 | 1.995 |
| 6MY2 | 1.399 | 1.126 | 3LO6 | 0.697 | 0.717 |
| 1OMC | 1.032 | 1.864 | 7N21 | 1.513 | 1.751 |
| 6MY1 | 0.688 | 1.388 | 7N24 | 1.396 | 1.892 |
| 2M62 | 2.646 | 2.295 | 7N25 | 1.026 | 2.831 |
| 5UG3 | 2.116 | 2.714 | 7N20 | 1.728 | 2.760 |
| 1KFP | 1.241 | 1.703 |  |  |  |
| **Mean** | 1.455 | 1.818 |  |  |  |
| **Median** | 1.396 | 1.703 |  |  |  |

Figure 2a compares two RMSD metrics between RFAA (baseline model) and NCPepFold. For cyclic peptide monomers, the average peptide RMSD decreased from 4.327 Å to 1.640 Å, while the median dropped from 4.008 Å to 1.609 Å. Similarly, for cyclic peptides with disulfide bonds, the average peptide RMSD decreased from 1.724 Å to 1.454 Å, and the median from 1.494 Å to 1.396 Å. The modification RMSD were slightly lower than the peptide RMSD values. Specifically, the modification RMSD for cyclic peptide monomers was 1.562 Å, and for disulfide-bonded cyclic peptides, it was 1.818 Å, representing performance improvements of 45.2% and 17.4%, respectively, compared to RFAA. Figure 2f shows representative predicted structures of two variants of θ-Defensins (PDB: 2M1P, 2M2G).^[37]^ θ-Defensins are ribosomally synthesized cyclic peptides found in the leukocytes of certain primates, featuring a β-sheet secondary structure. They hold significant potential as antimicrobial agents and scaffolds for peptide-based drugs. The structures predicted by NCPepFold are highly consistent with ground truth, particularly for 2M1P, where the peptide RMSD reached 1.239 Å. These results confirm that the assembly-based approach of NCPepFold enables accurate structure prediction for peptides with modified residues.

**Figure 2.**
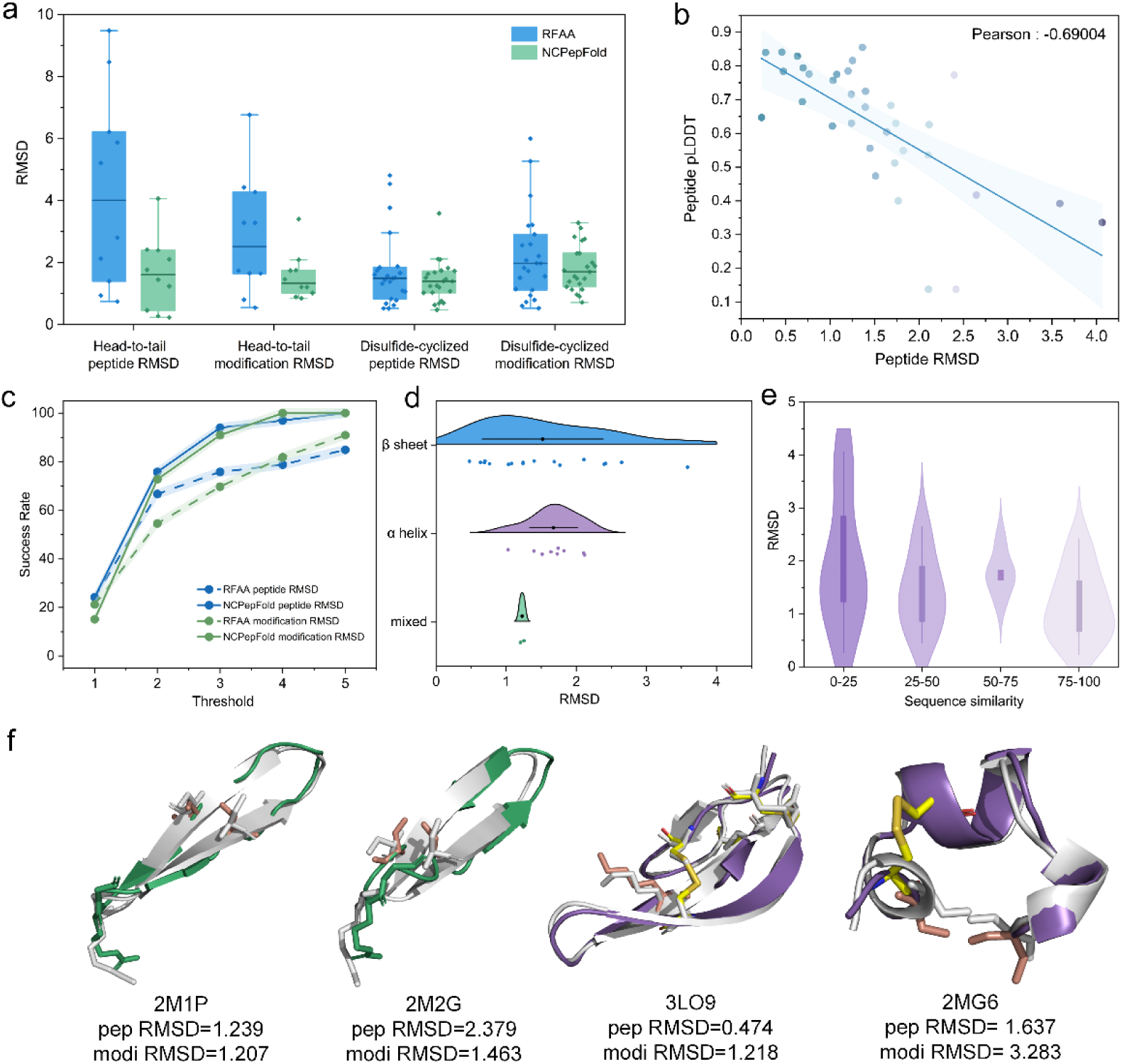
Predictive performance of NCPepFold on cyclic peptide structure prediction tasks. a) Peptide RMSD and modification RMSD metrics for head-to-tail cyclized and disulfide-bonded cyclic peptide monomers. Blue indicates results from RFAA, while green indicates results from NCPepFold. b) Correlation between peptide RMSD and peptide pLDDT. Pearson correlation coefficient is −0.69, which represents a strong negative correlation. c) SR of NCPepFold at different RMSD thresholds. Blue represents peptide RMSD, green represents modification RMSD, solid lines represent SR of NCPepFold, and dashed lines represent SR of RFAA. d) Relationship between different secondary structures and model accuracy. e) Comparison of sequence identity with the training set and model accuracy. The model is generally accurate even for sequences with low homology to the training set. f) Examples of cyclic peptide structures predicted by NCPepFold. Green and purple represent the predicted structures by NCPepFold, while gray represents the ground truth structure. (PDB IDs: 2M1P, 2M2G, 3LO9, 2MG6; where 2M1P and 2M2G are head-to-tail cyclic peptides, and 3LO9 and 2MG6 are disulfide-bonded cyclic peptides)

We report accuracy in terms of the percentage of successful predictions. Figure 2c illustrates success rates (SR) at different thresholds. SR is defined as the percentage of examples with RMSD below a given threshold. NCPepFold significantly improved SR under both strict (RMSD < 2 Å) and relaxed (RMSD < 5 Å) criteria. Specifically, under the strict criterion, NCPepFold achieved an SR of 75.8%, representing an approximate 9.1% improvement over RFAA. These observations collectively indicate that NCPepFold provides a model representation better suited for peptide structure prediction.

In Figure 2b, we observed a strong correlation between peptide RMSD and peptide pLDDT, with a Pearson correlation coefficient of −0.69. Notably, 77% of the predicted structures with a peptide RMSD < 2 Å can be accurately identified using peptide pLDDT, with a low false positive rate of 15.4%. Namely, pLDDT is a reliable metric to evaluate the quality of predicted structure and can be served as an effective indicator to filter potential compounds when experimental resources are limited.

Furthermore, we demonstrated the robustness of NCPepFold in terms of generalization. We binned the test set based on sequence similarity and calculated peptide RMSD for targets with varying levels of difficulty. Data with high structural similarity to the training set (similarity > 75%) were categorized as “simple,” those with moderate similarity (50% < similarity < 75%) as “moderate,” low similarity (25% < similarity < 50%) as “difficult,” and very low similarity (similarity < 25%) as “extremely difficult.” Across different levels of difficulty, NCPepFold showed improved predictive performance as sequence identity increased (Figure 2e). When we further examined more challenging targets, it was evident that NCPepFold still exhibited strong adaptability when dealing with unfamiliar scenarios. In difficult mode, the average peptide RMSD of the predicted structures reached 1.959 Å. By incorporating atom-level information about the target sequences and fine-tuning the model, NCPepFold effectively learned peptide-chain interaction information, potentially leveraging generalization capabilities beyond known protein conformations to achieve higher folding accuracy.

### 4.2. Accurate prediction of linear peptide structure

We added a cyclization option in NCPepFold, extending model’s functionality to linear peptides. We evaluated the performance of NCPepFold on 18 linear peptide monomers and 22 linear peptide complexes containing non-canonical amino acids. Figure 3a and 3d show the significant similarity between the predicted structures and the experimental structures for both peptide monomers and complexes. For monomers, the mean peptide RMSD was 1.646 Å, and the mean modification RMSD was 2.289 Å. Compared to RFAA, the peptide RMSD and modification RMSD for monomers improved by 40% and 16.3%, respectively.

**Figure 3.**
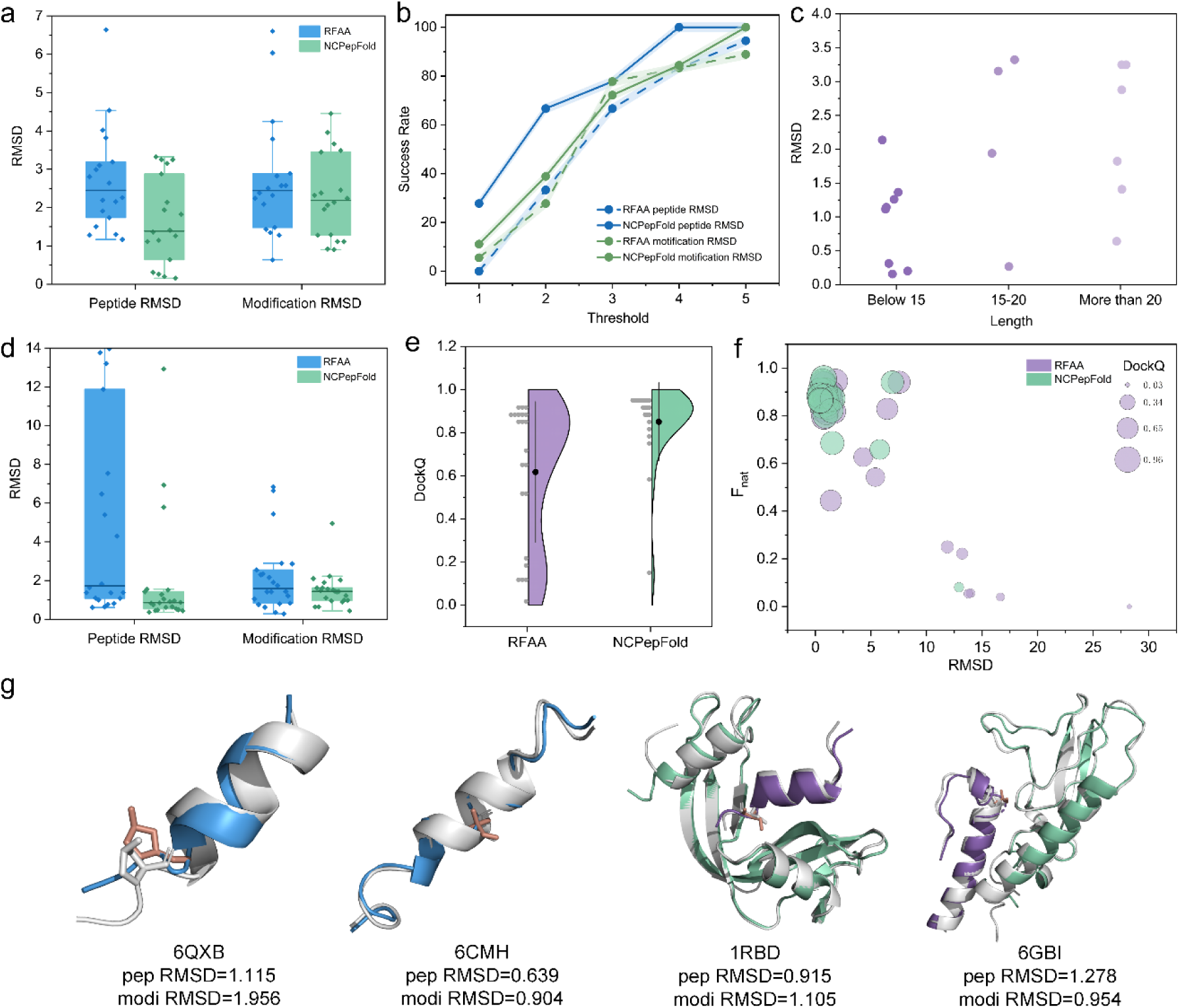
Predictive performance of NCPepFold on linear peptide structure prediction tasks. a) Peptide RMSD and modification RMSD metrics for linear peptide monomers. b) SR of NCPepFold at different RMSD thresholds. c) Relationship between peptide length and model accuracy. d) Peptide RMSD and modification RMSD metrics for linear peptide complexes. e) DockQ metrics for linear peptide complexes. f) Correlation among RMSD, Fnat, and DockQ. (G) Examples of linear peptide structures predicted by NCPepFold. Blue and purple represent the predicted structures by NCPepFold, while gray represents the ground truth structure. (PDB IDs: 6QXB, 6CMH, 1RBD, 6GBI; where 6QXB and 6CMH are linear peptide monomers, and 1RBD and 6GBI are linear peptide complexes)

**Figure 4.**
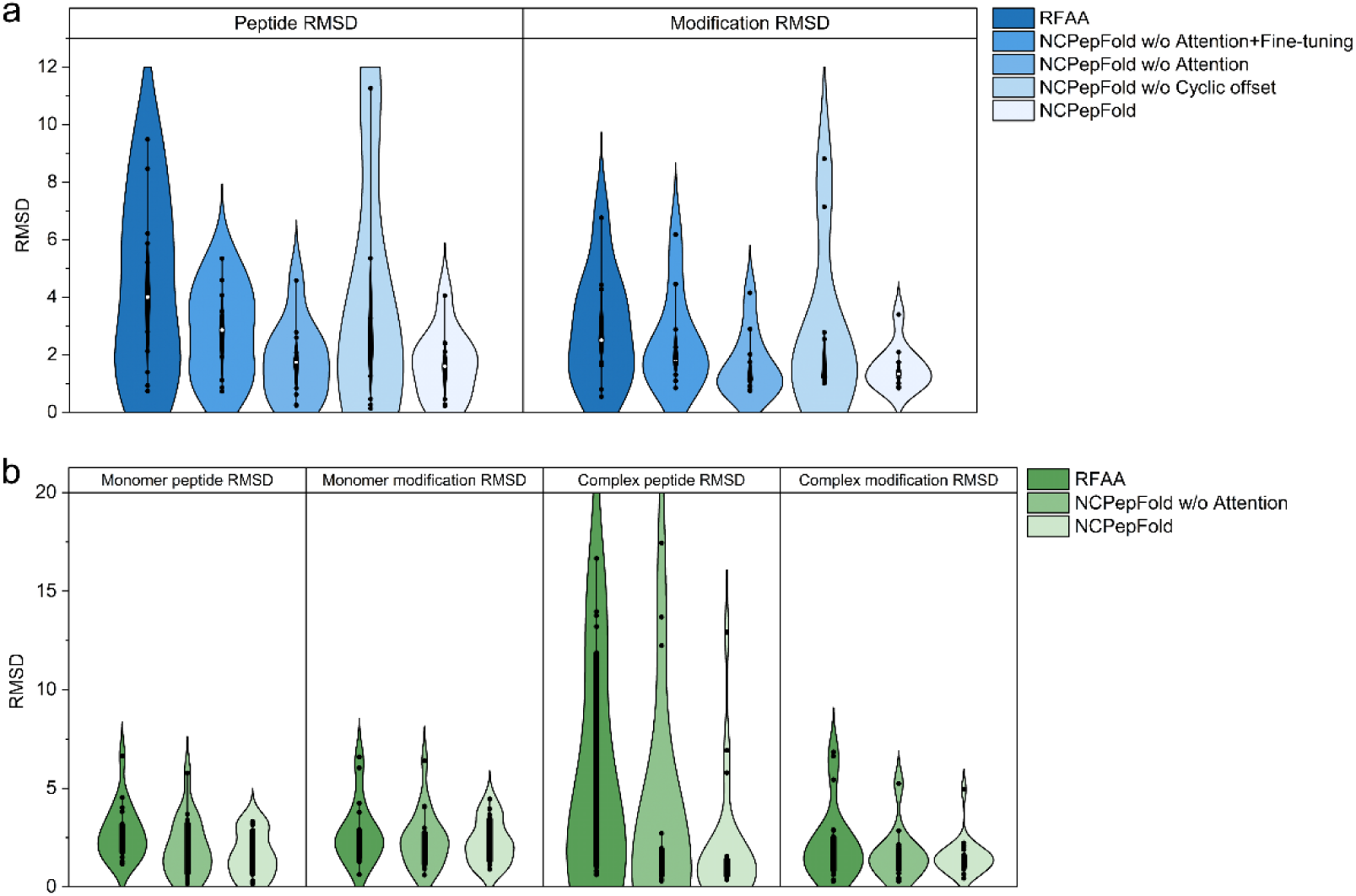
a) Ablation experiments results of cyclic peptides. b) Ablation experiments results of linear peptides.

In linear peptide-protein docking, NCPepFold demonstrated impressive effectiveness (**Table 4**). The mean peptide RMSD for complexes was 1.894 Å, while the mean modification RMSD was 1.509 Å. Notably, RFAA’s average peptide RMSD for predicting complex structures was 6.137 Å. We speculate that this is primarily because complex prediction requires not only the accurate reconstruction of the peptide chain conformation but also precise localization of the binding site on the target protein. Unlike monomer structure prediction, complex prediction is inherently more challenging, as any error in selecting the binding site significantly impacts the spatial configuration of the entire complex. Even if the peptide chain itself is accurately reconstructed, substantial deviations in binding site positioning prevent the final complex structure from closely approximating the experimental truth. Hence, the high RMSD observed in RFAA’s complex predictions mainly reflects its limitations in identifying and positioning the binding site, rather than merely errors in peptide chain conformation prediction. In contrast, NCPepFold outperformed RFAA significantly in complex prediction. DockQ was used to assess the accuracy of complex docking, with DockQ > 0.8 indicating a high-quality prediction. The median DockQ score for complexes predicted by NCPepFold was 0.911 (Figure 3e), a notable improvement over RFAA (0.783). The increase in DockQ scores indicates an enhancement in both binding site identification and the overall quality of the predicted complex structures. The optimized model not only achieved DockQ scores closer to ideal values but also reduced variability in docking accuracy. The success rate metric further substantiated this improvement in predictive accuracy (Figure 3b).

**Table 4.** Results of linear peptide monomer and complex.

| Monomer |  |  | Complex |  |  |  |  |
| --- | --- | --- | --- | --- | --- | --- | --- |
| PDB ID | Peptide RMSD | Modification RMSD | PDB ID | Peptide RMSD | Modification RMSD | DockQ | F<sub>nat</sub> |
| 2L87 | 1.94 | 4.455 | 1D5E | 1.048 | 1.518 | 0.903 | 0.850 |
| 2MZA | 3.156 | 2.139 | 6GB1 | 1.278 | 0.954 | 0.771 | 0.892 |
| 2N0N | 1.143 | 3.662 | 3OQZ | 0.619 | 1.451 | 0.881 | 0.841 |
| 2LDA | 2.137 | 2.061 | 2FX8 | 12.923 | 1.422 | 0.139 | 0.081 |
| 3ZS2 | 3.25 | 2.458 | 2K7L | 1.513 | 1.565 | 0.807 | 0.686 |
| 5KGY | 1.408 | 3.96 | 2RLN | 0.477 | 1.436 | 0.951 | 0.907 |
| 2LDD | 1.364 | 2.38 | 1RBD | 0.915 | 1.105 | 0.923 | 0.884 |
| 1GEA | 3.25 | 1.292 | 1D5D | 0.450 | 2.029 | 0.949 | 0.886 |
| 2LDC | 1.26 | 3.493 | 4O37 | 0.644 | 1.903 | 0.936 | 0.944 |
| 6QXB | 1.115 | 1.956 | 1FEV | 0.749 | 0.894 | 0.959 | 0.957 |
| 2MYL | 1.823 | 2.243 | 1D5H | 0.951 | 1.624 | 0.935 | 0.864 |
| 2MZ2 | 3.323 | 3.451 | 3KMZ | 0.833 | 0.989 | 0.894 | 0.882 |
| 2MYM | 2.878 | 2.321 | 4OKF | 0.539 | 1.480 | 0.935 | 0.864 |
| 2NOR | 0.2 | 1.116 | 1Z3P | 0.368 | 1.224 | 0.951 | 0.884 |
| 1V50 | 0.266 | 1.282 | 1Z3L | 0.495 | 0.644 | 0.936 | 0.861 |
| 6CMH | 0.639 | 0.904 | 4YJW | 1.557 | 0.978 | 0.902 | 0.872 |
| 3CMH | 0.311 | 0.917 | 4LKA | 0.878 | 1.623 | 0.882 | 0.800 |
| 1MXQ | 0.156 | 1.114 | 3OQY | 1.421 | 0.440 | 0.847 | 0.822 |
|  |  |  | 1Z3M | 0.442 | 0.639 | 0.942 | 0.865 |
|  |  |  | 5OTT | 5.789 | 4.947 | 0.582 | 0.659 |
|  |  |  | 5OTU | 6.928 | 2.105 | 0.763 | 0.941 |
|  |  |  | 3OR0 | 0.849 | 2.227 | 0.918 | 0.907 |
| **Mean** | 1.646 | 2.289 |  | 1.894 | 1.509 | 0.850 | 0.825 |
| **Median** | 1.386 | 2.191 |  | 0.864 | 1.444 | 0.911 | 0.869 |

Figure 3f shows representative predicted structures of two protein-peptide complexes: ribonuclease S (RNase-S, PDB: 1RBD) and the GLP-1 receptor with a bound peptide (PDB: 6GBI).^[38,39]^ Bovine pancreatic ribonuclease A (RNase-A) can be hydrolyzed by trypsin into two fragments, which can reassemble to form catalytically active ribonuclease S. The predicted structure of RNase-A by NCPepFold achieved a peptide RMSD of 0.915 and a DockQ score of 0.923, demonstrating high consistency with the experimental structure.

### 4.3. Ablation experiments

We conducted a series of ablation experiments using RFAA as the baseline model to explore the impact of fine-tuning strategies and individual modules on NCPepFold. Notably, cyclic peptides incorporate an additional cyclization offset module compared to linear peptides, while all other modules remain identical. As shown in **Table 5** and **Table 6**, the original version of NCPepFold demonstrated the best performance across all architectures.

**Table 5.** Ablation experiments results of cyclic peptides.

| Methods | Peptide RMSD ↓ | Modification RMSD ↓ |
| --- | --- | --- |
| RFAA (baseline) | 4.327 | 2.842 |
| NCPEPfold w/o Attention + Fine-tuning | 2.792 | 2.433 |
| NCPEPfold w/o Attention | 1.753 | 1.649 |
| NCPEPfold w/o CPM | 3.777 | 2.816 |
| NCPEPfold | **1.640** | **1.562** |

**Table 6.** Ablation experiments results of linear peptides.

| Methods | Peptide RMSD↓ | Modification RMSD↓ | DockQ↑ | F<sub>nat</sub>↑ |
| --- | --- | --- | --- | --- |
| RFAA (baseline) | 4.607 | 2.428 | 0.617 | 0.616 |
| NC PepFold w/o attention | 3.243 | 1.989 | 0.771 | 0.760 |
| NC PepFold | **1.782** | **1.860** | **0.850** | **0.825** |

When Attention module was removed from NCPepFold, the model’s performance declined in both peptide RMSD and modified RMSD metrics. Specifically, the peptide RMSD for cyclic peptides increased from 1.64 Å to 1.753 Å, and for linear peptides, it rose from 1.782 Å to 3.243 Å. This indicates that the Attention module effectively captures long-range interactions between residues and atomic interactions during the feature extraction process. Representing these interactions is critical for improving the overall accuracy of folding conformation prediction.

To investigate the impact of fine-tuning on model performance, we compared various metrics between the baseline model and NCPepFold w/o Attention for linear peptides, as well as between NCPepFold w/o Attention and NCPepFold w/o Attention + Fine-tuning for cyclic peptides. The results show that in the absence of peptide-specific structural information, the model exhibits performance degradation across all metrics to varying degrees. Specifically, the average peptide RMSD increased by 29.6% for linear peptides and 37.2% for cyclic peptides, while the modification RMSD increased by 18.1% and 32.2%, respectively. We hypothesize that peptides and proteins share domain relevance, with similarities in local structural patterns and dynamic properties. Fine-tuning enables the sharing of low-level features and the adaptation of high-level features, thereby further improving model accuracy.

CPM module is another critical component in NCPepFold specifically designed for cyclic peptides. Removing CPM resulted in a significant performance decline, with the peptide RMSD increasing from 1.640 Å to 3.777 Å and the modification RMSD increasing from 1.562 Å to 2.816 Å. These observed performance degradations underscore the essential role of CPM module. CPM introduces a soft constraint on the terminal residues, greatly enhancing the likelihood of forming a cyclic structure. Additionally, integrating CPM with baseline module reduced the peptide RMSD from 4.327 Å to 2.792 Å, further validating its effectiveness.

## 5. Conclusion

Although the importance of nonstandard amino acids for the chemogenic application of peptides has long been recognized, how to represent nonstandard amino acids would have been difficult before the recent advances in deep learning–based peptide structure prediction. In this study, we developed NCPepFold, an approach built upon RFAA that integrates a cyclic position matrix algorithm with the Transformer framework and employs transfer learning on various types of peptide data. This makes it a specialized model for predicting the structures of cyclic peptides with non-canonical amino acids. As an end-to-end deep learning approach, NCPepFold fills the gap in structure prediction of cyclic peptide with non-canonical amino acids. Unlike traditional single-representation methods for residues, NCPepFold achieves a seamless integration of residue representation with atomic graph representation. This multi-level information integration enhances peptide representation capabilities, elevating the prediction accuracy of non-canonical amino acid-containing peptides to the atomic level. We tested 73 peptides across five groups, demonstrating that NCPepFold shows a clear advantage in secondary structures predictions of both cyclic and linear peptides. Overall, except for a few outliers, the peptide structures were predicted with good success. In monomers, deviations caused by these outliers were primarily due to errors in the loop regions, which are inherently flexible and may result in multiple conformations for the same structure. For complexes, the outliers were influenced not only by structural deviations in the loop regions but also by incorrect identification of binding sites.

Notably, NCPepFold is the first model capable of specifically predicting the structure of cyclic peptides with non-canonical amino acids with sequence input alone. The peptide structures generated by NCPepFold may provide valuable insights into the inference of non-canonical amino acids towards local conformations of peptides, potentially clarifying structure-function relationships and augmenting our mechanistic understanding.

During our work, AlphaFold3 (AF3) was open-sourced and demonstrated high accuracy in predicting nearly all types of molecules found in PDB, including chemically modified proteins.^[40]^ However, AF3 lacks specific constraints for cyclic structures and heavily relies on templates from sequence alignment. In other words, if the template is linear, AF3 will tend to predict a linear structure, which significantly limits its ability to predict cyclic peptides. Notably, cyclic peptides are typically first synthesized as linear peptides using solid-phase peptide synthesis (SPPS), and then cyclized through chemical or enzymatic methods.^[41]^ This means cyclization is a controlled process determined by experimental design, so whether a peptide is cyclic or linear depends on researcher’s needs. While AF3 performs excellently in linear peptide structure prediction, it still falls short in cyclic structures. To overcome this limitation, we may consider integrating cyclization constraints into AF3 in the future to improve prediction accuracy for cyclic peptides and proteins with complex spatial structures.

However, our model also has some limitations. Firstly, the training data may not cover all non-canonical amino acids, especially rare ones, leading to less accurate predictions for the side-chain conformations of certain amino acids with chemical modification. Secondly, predicting the cyclization of a small molecule linking two residues on a same peptide chain to form a ring is still a significant challenge. Finally, for cases where the binding sites of some complexes were not correctly identified, we believe that adding binding site information to the model could substantially enhance the prediction accuracy of the complexes. Moving forward, our research will focus on improving the model to address these issues and further advance the field of structural biology.

## Supporting Information

Supporting Information is available from the Wiley Online Library or from the author. sample

## Acknowledgements

This project was supported by the Natural Science Foundation of Zhejiang Province (LD22H300004).

## Author contributions

Q. M., T. S.: These authors contributed equally to this work.

Conceptualization and investigation: Q. M., T. S., H. D. Data acquisition: Q. M., W. X. Develop the model: Q.M., T.S., S.Z. Visualized data and created figures: Q. M., W.X. Writing-original draft: Q. M. Writing-review & editing: Q. M., T. S., C. Z. Funding acquisition: H. D. Project administration: C. L., A. S.

## Data Availability Statement

The data that support the findings of this study are available from the corresponding author upon reasonable request.

