## Supporting Information for "NCPepFold: Accurate Prediction of Non-canonical Cyclic Peptide Structures via Cyclization Optimization with Multigranular Representation"

**Section S1. Datasets**

***Training datasets***

*Head-to-tail peptide dataset*

The cyclic peptide dataset comes from the work of HighFold, an AI model for predicting cyclic peptide structures, including both monomers and complexes.^[1]^ The cyclic peptide monomers dataset, obtained from the work of Bhardwaj, comprises 63 experimentally investigated structures ranging from 12 to 39 residues.^[2]^ These structures encompass 7 α-helical and 37 β-sheet, 10 random coil and 9 mixed secondary structure motifs. The cyclic peptide monomers dataset, derived from the work of ADCP, comprises 17 structures with ligand lengths spanning 6–14 residues.^[3]^ Secondary structure motifs within the ligands include 3 β-sheet, 14 random coil structures and 2 mixed structures.

The PDB IDs of the HighFold dataset are as follows:

Monomer: 1BH4, 1DF6, 1HVZ, 1JBL, 1QVK, 1QVL, 1R1F, 1SKI, 1SKK, 1VB8, 1ZA8, 1ZNU, 2ATG, 2B38, 2ERI, 1GJ0, 2K7G, 2KCH, 2KNM, 2KUK, 2KUX, 2KVX, 2LAM, 2LL5, 2LUR, 2LWS, 2LWT, 2LWU, 2LWV, 2LYE, 2LYF, 2LZI, 2M77, 2M78, 2M79, 2M9O, 2MH1, 2MN1, 2MSO, 2MT8, 2MW0, 2N07, 2NB5, 2NDL, 2NDM, 2NDN, 2NS4, 2OTQ, 2OX2, 2PO8, 5H1H, 5H1I, 5KWZ, 5KX1, 5WOV, 5WOW, 6DNY, 6PIN, 6PIO, 6PIP, 6U7Q, 6U7R, 6U7S, 6WPV, 7F32, 7K7X, 7I53, 7I54, 7I55, 7IHC, 7M25, 7M27, 7M28, 7M29, 7M2A, 7M2B, 7M2C, 7M3U, 7RN3, 7S55

Complex: 1SFI, 3AV9, 3AVA, 3AVB, 3AVF, 3AVG, 3AVH, 3AVI, 3AVJ, 3AVK, 3AVM, 3AVN, 3P8F, 3WNE, 3ZGC, 4K1E, 4KEL

*Disulfide-Cyclized Peptide Dataset*

The disulfide-cyclized peptide dataset includes 46 monomers and 62 complexes. The monomers are sourced from CyclicPepedia, while the complexes are from Propedia.^[4,5]^ Propedia is a protein-peptide interaction database that uses the following criteria to retrieve PDB entries: (1) structures composed by two or more chains, (2) one chain with at least 2 and no more than 50 residues (for peptides), and (3) structures solved by NMR or X-ray crystallography with resolution below 2.5 Å. The present release is composed of 19,813 complexes (May 02, 2020). We extracted structures containing two or more cysteine residues (CYS), ultimately obtaining 62 disulfide-cyclized peptide complexes.

The PDB IDs of monomers are as follows: 1CZ6, 1DFN, 1EWS, 1F3K, 1FEO, 1FU3, 1IXT, 1JBN, 1LFC, 1MA2, 1N1U, 1RMK, 1RPB, 1RPC, 1S6W, 1TV0, 1X7K, 1ZMQ, 2EFZ, 2FQA, 2IFJ, 2IGU, 2JNI, 2JSB, 2JUQ, 2JUR, 2JUS, 2K10, 2K1I, 2KM9, 2LXZ, 2M3N, 2MD6, 2MSO, 2N7F, 5XO3, 5XO4, 5XO5, 5Y0H, 6CEI, 6EFE, 6OTB, 6Q5Z, 7N0T, 7N23, 7RFA.

The PDB IDs of complexes are as follows: 1F2S, 1G9I, 1GL0, 1H9I, 1HQQ, 1HXZ, 1JBU, 1JDP, 1MCT, 1OX1, 1PMX, 1PPE, 1SBW, 2BTC, 2CK0, 2NWN, 2O9Q, 2PLX, 2STA, 2XTT, 3G5V, 3G5Y, 3IIQ, 3M61, 3OX7, 3OY5, 3OY6, 3P72, 3PP4, 3SRI, 3ZDF, 4AOQ, 4GW1, 4GW5, 4IB5, 4U2W, 4W50, 4X1N, 4X1Q, 4X1R, 4XOJ, 4Z09, 4Z0D, 5DI8, 5DJ0, 5DVL, 5EOC, 5EUK, 5F88, 5FF6, 5H5S, 5JZU, 5TH2, 5WN9, 6A8N, 6GD5, 6IGK, 6Q9F, 7EGR, 7T10, 7WIC, 7WJ5.

*ncAA Linear Peptide Dataset*

The linear peptide dataset with non-canonical amino acids was extracted from the test set provided on PEPstrMOD developers' website.^[6]^ To evaluate PEPstrMOD's performance in handling modified peptides, first, workers extracted 21182 PDB chains having length between 7 and 25. Next, they searched for D-amino acids, non-natural amino acids as specified in FFNCAA and post-translational modifications as specified in FFPTM library and obtained 47, 72 and 692 (total 811) PDB chains. Further, they removed peptides having disulfide bridges and duplicate sequences. Finally, a dataset containing 501 PDB chains was obtained and named the ModPep dataset. ModPep includes 56 monomers and 445 complexes. We first replaced the non-canonical amino acids in the 501 entries with their standard amino acid analogs, and then used RFAA to predict the structures of these entries. Structure generation failed for 282 samples due to three main reasons: 1. CUDA out of memory; 2. No such file or directory: 'data/…/t000.atab'; 3. need at least one array to concatenate. The first reason was insufficient memory caused by excessively long protein sequences. The second was missing files due to issues with multiple sequence alignment (MSA). The third occurred when the number of repeated residues between the target sequence and its similar sequences was fewer than 10, resulting in an empty list being passed during the initialization of coordinates, leading to this error. From the remaining usable data, we filtered out structures containing D-amino acids, structures with breaks in the peptide chain, structures where atoms in non-canonical amino acids were missing, and 1 cyclic peptide structure (2M2X). Finally, we obtained 37 monomers and 181 complexes. Additionally, we extracted samples with a secondary structure content greater than 50% to form the test set. In the end, we obtained 32 linear peptide monomers and 174 linear peptide complexes as part of the training set.

The PDB IDs of monomers are as follows: 1T9E, 2CEF, 2CEZ, 2CFJ, 2D3H, 2D3H, 2D3H, 2D3H, 2D3H, 2DPR, 2GFR, 2JQC, 2L5J, 2OYV, 3A08, 3A08, 3A08, 3A08, 3A0A, 3A0A, 3A0A, 3A0A, 3A0A, 3A0A, 3A0M, 3A0M, 3A0M, 3A0M, 3A1H, 3A1H, 3A1H, 3ABN.

The PDB IDs of complexes are as follows: 1AI1, 1AOT, 1CFN, 1F58, 1FGL, 1FPR, 1FU5, 1G6G, 1GUW, 1I8G, 1I8H, 1IRS, 1J4P, 1J4Q, 1JLU, 1JM4, 1JSP, 1JU5, 1LM8, 1LQB, 1MW4, 1O9S, 1P16, 1P22, 1PFB, 1PUA, 1Q1A, 1QG1, 1SHC, 1SPS, 1T29, 1T2V, 1TCE, 1UEF, 1YRK, 2A0T, 2BQZ, 2CI9, 2DVQ, 2DVR, 2E3K, 2FUU, 2G46, 2HMH, 2IUI, 2JND, 2JQI, 2JQL, 2K17, 2KTB, 2KVM, 2KWN, 2L11, 2L1B, 2L3R, 2LAJ, 2LB0, 2LB2, 2LCT, 2LGK, 2LO6, 2LSP, 2LUE, 2LVM, 2LYW, 2M0O, 2M3M, 2MC1, 2MFQ, 2MJV, 2OQ1, 2OS2, 2OVQ, 2P5B, 2PLD, 2PNX, 2RMX, 2RNW, 2RNX, 2RNY, 2ROR, 2RR4, 2RSN, 2V87, 2V89, 2WP1, 2WP2, 2X4W, 2X4X, 2Y5T, 2YBP, 2YU7, 3AL3, 3ASK, 3AVR, 3BH8, 3BH9, 3BHB, 3BU6, 3BUO, 3COJ, 3DM1, 3DPC, 3DPP, 3DPQ, 3EJH, 3FQR, 3HNA, 3I90, 3KV4, 3L6F, 3MHR, 3N9N, 3N9P, 3O34, 3OB1, 3OB2, 3OP0, 3PDH, 3PFV, 3PQZ, 3QO2, 3R93, 3SQD, 3SVM, 3TU7, 3TZD, 3U5N, 3UVW, 3UVY, 3WDZ, 3WP0, 3WP1, 3ZKF, 4AJY, 4APJ, 4BJ3, 4BKL, 4BL0, 4BU0, 4BU1, 4DOW, 4EZH, 4GAG, 4GLR, 4HCZ, 4HON, 4HPT, 4I7B, 4JXT, 4L1U, 4LLB, 4LOR, 4N4I.

***Test datasets***

Test set includes 18 linear peptide monomers, 22 linear peptide complexes, 10 head-to-tail cyclic peptide monomers, and 23 disulfide-cyclized peptide monomers.

We used DSSP to filter linear peptide monomers and complexes with more than 50% secondary structure content in ncAA Linear Peptide Dataset.^[7]^ The DSSP settings are as follows: We used DSSP software to assign secondary structure states to the peptide residues. DSSP defines eight states (T, S, G, H, I, B, E, −), where 'H' is considered α-helix, 'B' is considered β-sheet, and states 'S' and 'C' are classified as loops. This process resulted in 6 linear peptide monomers and 9 linear peptide complexes. Additionally, we manually selected 12 linear peptide monomers and 13 linear peptide complexes from the PDB database, which, together with the previous data, formed the linear peptide test set. Among the 18 linear peptide monomers, 11 contained one non-canonical amino acid, while 7 contained two non-canonical amino acids, and all adopted α-helical conformations. This is because β-sheets require at least two interacting strands to form a stable structure, and short linear peptides generally lack sufficient residues for such interactions. For the 22 linear peptide complexes, 15 had one non-standard amino acid, 6 had two, and 1 had three, all of which were α-helical. This is because there were virtually no linear peptide complexes with β-sheets that met the secondary structure requirements.

The PDB IDs of the linear peptide monomers in the ModPep dataset are as follows: 1MXQ, 1V50, 2NOR, 3CMH, 3ZS2, 6CMH.

The PDB IDs of the linear peptide monomers selected from the PDB database are as follows: 1GEA, 2LDA, 2LDD, 2LDC, 6QXB, 2MYL, 2MYM, 2MZ2, 2MZA, 2N0N, 5KGY, 2L87.

The PDB IDs of the linear peptide complexes in the ModPep dataset are as follows: 1FEV, 1RBD, 1Z3L, 1Z3M, 1Z3P, 2FX8, 2K7L, 3KMZ, 4LKA.

The PDB IDs of the linear peptide monomers selected from the PDB database are as follows: 1D5D, 1D5H, 1D5E, 2RLN, 3OQZ, 3OR0, 3OQY, 4OKF, 4O37, 4YJW, 5OTT, 5OTU, 6GB1.

10 head-to-tail cyclic monomers were sourced from CyclicPepedia, CPPsite, and additional similar structures found in the PDB database using the 'Find Similar Assemblies' option.^[4,8]^ Among these, 5 contain one non-canonical amino acids, 3 contain two non-canonical amino acids, 1 contains four non-canonical amino acids, and 1 contains six non-canonical amino acids, all of which adopt β-sheet conformations. Due to the limited number of available data, we relaxed the search criteria to require only the presence of secondary structures. The selection criteria from CyclicPepedia and CPPsite were as follows: the peptide chain must have secondary structures, contain 10 or more residues, with non-canonical amino acids comprising no more than 30% of the total, and no D-amino acids present.

The PDB IDs of the head-to-tail cyclic peptide monomers are as follows: 1T9E, 2J15, 2KNN, 2M1P, 2M2G, 2M2S, 2M2X, 2MSQ, 2N8E, 4M6E.

Among 23 disulfide-cyclized peptide monomers, 20 contain one non-canonical amino acids, 2 contain two non-canonical amino acids, 1 contain three non-canonical amino acids, with 12 adopting β-sheet structures, 9 adopting α-helical structures, and 2 having a mixed α-helix and β-sheet structure.

The PDB IDs of the β-sheet structures are as follows: 1KFP, 1OMC, 2EW4, 2M62, 3HJD, 3LO6, 3LO9, 4E83, 4E86, 6MY1, 6MY2, 6MY3.

The PDB IDs of the α-helix structures are as follows: 1K64, 2MG6, 2MFX, 5UG3, 7N21, 7N25, 7N20, 7N24, 1P9G.

The PDB IDs of the mixed structures are as follows: 1BIG, 2CRD.

**Section S2.** Results of peptide RMSD and modification RMSD

| **PDB ID** | **RFAA** | **NCPepFold w/o Attention** | **NCPepFold** |
| --- | --- | --- | --- |
| 2L87 | 2.639 | 1.912 | 1.940 |
| 2MZA | 1.739 | 3.687 | 3.156 |
| 2N0N | 3.818 | 0.988 | 1.143 |
| 2LDA | 2.194 | 1.838 | 2.137 |
| 3ZS2 | 4.534 | 3.260 | 3.250 |
| 5KGY | 1.912 | 1.553 | 1.408 |
| 2LDD | 1.299 | 1.294 | 1.364 |
| 1GEA | 3.101 | 3.260 | 3.250 |
| 2LDC | 2.156 | 0.677 | 1.260 |
| 6QXB | 1.503 | 1.709 | 1.115 |
| 2MYL | 1.169 | 1.836 | 1.823 |
| 2MZ2 | 1.280 | 2.489 | 3.323 |
| 2MYM | 2.991 | 3.402 | 2.878 |
| 2NOR | 3.189 | 0.241 | 0.20 |
| 1V50 | 2.260 | 0.538 | 0.266 |
| 6CMH | 6.640 | 5.775 | 0.639 |
| 3CMH | 2.802 | 0.431 | 0.311 |
| 1MXQ | 4.022 | 0.169 | 0.156 |

**Table S1.** Results of peptide RMSD for linear peptide monomers

| **PDB ID** | **RFAA** | **NCPepFold w/o Attention** | **NCPepFold** |
| --- | --- | --- | --- |
| 2L87 | 6.599 | 6.396 | 4.455 |
| 2MZA | 1.480 | 2.570 | 2.139 |
| 2N0N | 6.034 | 2.990 | 3.662 |
| 2LDA | 2.317 | 2.329 | 2.061 |
| 3ZS2 | 2.504 | 1.818 | 2.458 |
| 5KGY | 3.788 | 4.054 | 3.960 |
| 2LDD | 2.824 | 2.448 | 2.380 |
| 1GEA | 2.087 | 1.141 | 1.292 |
| 2LDC | 2.570 | 2.505 | 3.493 |
| 6QXB | 1.346 | 2.328 | 1.956 |
| 2MYL | 2.385 | 2.212 | 2.243 |
| 2MZ2 | 4.247 | 4.092 | 3.451 |
| 2MYM | 2.575 | 2.782 | 2.321 |
| 2NOR | 2.244 | 0.932 | 1.116 |
| 1V50 | 1.282 | 1.508 | 1.282 |
| 6CMH | 1.434 | 0.612 | 0.904 |
| 3CMH | 0.638 | 0.918 | 0.917 |
| 1MXQ | 2.891 | 1.059 | 1.114 |

**Table S2.** Results of modification RMSD for linear peptide monomers

| **PDB ID** | **RFAA** | **NCPepFold w/o Attention** | **NCPepFold** |
| --- | --- | --- | --- |
| 1D5E | 0.647 | 0.640 | 1.048 |
| 6GB1 | 1.818 | 1.321 | 1.278 |
| 3OQZ | 0.766 | 0.542 | 0.619 |
| 3OR0 | 1.004 | 0.643 | 0.849 |
| 2FX8 | 28.25 | 34.228 | 12.923 |
| 2K7L | 5.396 | 1.959 | 1.513 |
| 2RLN | 1.371 | 0.687 | 0.477 |
| 1RBD | 1.094 | 0.606 | 0.915 |
| 1D5D | 0.821 | 0.914 | 0.450 |
| 4O37 | 13.203 | 12.238 | 0.644 |
| 1FEV | 0.615 | 0.546 | 0.749 |
| 1D5H | 11.870 | 0.929 | 0.951 |
| 3KMZ | 4.296 | 2.719 | 0.833 |
| 4OKF | 1.635 | 0.365 | 0.539 |
| 1Z3P | 1.084 | 0.574 | 0.368 |
| 1Z3L | 13.964 | 0.296 | 0.495 |
| 4YJW | 1.373 | 1.594 | 1.557 |
| 4LKA | 16.655 | 0.703 | 0.878 |
| 3OQY | 1.388 | 1.625 | 1.421 |
| 1Z3M | 13.762 | 0.405 | 0.442 |
| 5OTT | 6.465 | 13.689 | 5.789 |
| 5OTU | 7.535 | 17.447 | 6.928 |

**Table S3.** Results of peptide RMSD for linear peptide complexes

| **PDB ID** | **RFAA** | **NCPepFold w/o Attention** | **NCPepFold** |
| --- | --- | --- | --- |
| 1D5E | 1.907 | 1.800 | 1.518 |
| 6GB1 | 1.417 | 0.681 | 0.954 |
| 3OQZ | 0.357 | 1.377 | 1.451 |
| 3OR0 | 2.163 | 1.963 | 2.227 |
| 2FX8 | 1.741 | 1.554 | 1.422 |
| 2K7L | 1.423 | 1.527 | 1.565 |
| 2RLN | 0.804 | 1.299 | 1.436 |
| 1RBD | 0.623 | 0.402 | 1.105 |
| 1D5D | 2.891 | 2.152 | 2.029 |
| 4O37 | 2.341 | 5.243 | 1.903 |
| 1FEV | 0.285 | 0.452 | 0.894 |
| 1D5H | 2.300 | 2.047 | 1.624 |
| 3KMZ | 1.435 | 1.644 | 0.989 |
| 4OKF | 0.761 | 1.306 | 1.480 |
| 1Z3P | 1.211 | 1.279 | 1.224 |
| 1Z3L | 2.549 | 0.390 | 0.644 |
| 4YJW | 0.831 | 0.840 | 0.978 |
| 4LKA | 5.435 | 1.085 | 1.623 |
| 3OQY | 1.055 | 1.404 | 0.440 |
| 1Z3M | 2.866 | 0.270 | 0.639 |
| 5OTT | 6.837 | 5.239 | 4.947 |
| 5OTU | 6.640 | 2.860 | 2.105 |

**Table S4.** Results of modification RMSD for linear peptide complexes.

| **PDB ID** | **RFAA** | **NCPepFold w/o Attention + Fine-tuning** | **NCPepFold w/o Attention** | **NCPepFold w/o Cyclic offset** | **NCPepFold** |
| --- | --- | --- | --- | --- | --- |
| 2N8E | 0.744 | 0.737 | 0.242 | 0.267 | 0.274 |
| 4M6E | 5.875 | 4.598 | 4.587 | 5.355 | 4.063 |
| 2J15 | 5.212 | 3.339 | 1.629 | 3.268 | 1.450 |
| 2MSQ | 0.937 | 0.863 | 0.627 | 0.469 | 0.458 |
| 2M2S | 6.217 | 4.071 | 2.789 | 11.263 | 1.767 |
| 2M1P | 1.402 | 1.123 | 0.846 | 1.269 | 1.239 |
| 2M2G | 2.804 | 2.396 | 1.860 | 2.288 | 2.397 |
| 2KNN | 2.129 | 3.512 | 2.078 | 2.183 | 2.108 |
| 2M2X | 9.483 | 5.346 | 2.600 | 11.263 | 2.418 |
| 1T9E | 8.471 | 1.937 | 0.267 | 0.141 | 0.229 |

**Table S5.** Results of peptide RMSD for head-to-tail cyclic peptide monomers.

| **PDB ID** | **RFAA** | **NCPepFold w/o Attention + Fine-tuning** | **NCPepFold w/o Attention** | **NCPepFold w/o Cyclic offset** | **NCPepFold** |
| --- | --- | --- | --- | --- | --- |
| 2N8E | 0.547 | 1.312 | 1.108 | 1.108 | 1.209 |
| 4M6E | 3.281 | 4.459 | 2.023 | 2.783 | 2.090 |
| 2J15 | 3.283 | 1.895 | 0.792 | 2.547 | 0.845 |
| 2MSQ | 1.653 | 1.098 | 0.751 | 1.147 | 0.895 |
| 2M2S | 4.275 | 2.885 | 2.901 | 7.147 | 1.753 |
| 2M1P | 0.801 | 0.847 | 0.902 | 1.210 | 1.207 |
| 2M2G | 1.741 | 1.714 | 1.176 | 1.212 | 1.463 |
| 2KNN | 1.645 | 1.661 | 1.751 | 1.178 | 1.741 |
| 2M2X | 6.765 | 6.184 | 4.156 | 8.816 | 3.402 |
| 1T9E | 4.426 | 2.273 | 0.925 | 1.015 | 1.019 |

**Table S6.** Results of modification RMSD for head-to-tail cyclic peptide monomer.

| **PDB ID** | **RFAA** | **NCPepFold w/o Attention** | **NCPepFold** |
| --- | --- | --- | --- |
| 1P9G | 4.538 | 4.122 | 2.104 |
| 2CRD | 4.812 | 1.087 | 1.203 |
| 1BIG | 1.382 | 1.171 | 1.252 |
| 1K64 | 1.571 | 1.934 | 1.740 |
| 2EW4 | 2.960 | 3.656 | 3.587 |
| 6MY3 | 1.494 | 2.066 | 1.684 |
| 6MY2 | 1.441 | 1.888 | 1.399 |
| 1OMC | 1.513 | 1.187 | 1.032 |
| 6MY1 | 1.322 | 1.957 | 0.688 |
| 2M62 | 3.769 | 1.896 | 2.646 |
| 5UG3 | 1.878 | 2.198 | 2.116 |
| 1KFP | 1.843 | 1.885 | 1.241 |
| 2MFX | 1.666 | 1.618 | 1.824 |
| 2MG6 | 0.828 | 1.883 | 1.637 |
| 3LO9 | 0.520 | 0.461 | 0.474 |
| 4E86 | 0.623 | 0.683 | 0.763 |
| 4E83 | 0.676 | 1.033 | 1.075 |
| 3HJD | 0.784 | 0.523 | 0.632 |
| 3LO6 | 0.523 | 0.561 | 0.697 |
| 7N21 | 1.743 | 1.757 | 1.513 |
| 7N24 | 1.089 | 1.888 | 1.396 |
| 7N25 | 1.609 | 1.168 | 1.026 |
| 7N20 | 1.063 | 1.315 | 1.728 |

**Table S7.** Results of peptide RMSD for disulfide-bonded cyclic monomers.

| **PDB ID** | **RFAA** | **NCPepFold w/o Attention** | **NCPepFold** |
| --- | --- | --- | --- |
| 1P9G | 1.118 | 2.471 | 2.301 |
| 2CRD | 2.591 | 0.925 | 1.313 |
| 1BIG | 1.729 | 1.184 | 1.490 |
| 1K64 | 1.982 | 1.653 | 1.132 |
| 2EW4 | 2.076 | 2.483 | 1.558 |
| 6MY3 | 0.941 | 1.659 | 1.484 |
| 6MY2 | 3.215 | 1.687 | 1.126 |
| 1OMC | 1.826 | 2.132 | 1.864 |
| 6MY1 | 1.975 | 1.695 | 1.388 |
| 2M62 | 2.908 | 0.583 | 2.295 |
| 5UG3 | 1.509 | 2.034 | 2.714 |
| 1KFP | 1.558 | 1.056 | 1.703 |
| 2MFX | 2.555 | 2.588 | 3.118 |
| 2MG6 | 2.210 | 3.574 | 3.283 |
| 3LO9 | 1.145 | 1.042 | 1.218 |
| 4E86 | 0.728 | 0.774 | 0.907 |
| 4E83 | 0.792 | 0.734 | 0.963 |
| 3HJD | 0.601 | 1.870 | 1.995 |
| 3LO6 | 0.525 | 0.600 | 0.717 |
| 7N21 | 5.270 | 3.185 | 1.751 |
| 7N24 | 4.160 | 3.542 | 1.892 |
| 7N25 | 3.192 | 1.576 | 2.831 |
| 7N20 | 6.000 | 4.114 | 2.760 |

**Table S8.** Results of modification RMSD for disulfide-bonded cyclic monomers.

Section S3. The evaluation index results of the complex

| **PDB ID** | **RFAA** | **NCPepFold w/o Attention** | **NCPepFold** |
| --- | --- | --- | --- |
| 1D5E | 0.859 | 0.931 | 0.903 |
| 6GB1 | 0.850 | 0.785 | 0.771 |
| 3OQZ | 0.888 | 0.889 | 0.881 |
| 3OR0 | 0.882 | 0.912 | 0.918 |
| 2FX8 | 0.032 | 0.024 | 0.139 |
| 2K7L | 0.512 | 0.720 | 0.807 |
| 2RLN | 0.867 | 0.934 | 0.951 |
| 1RBD | 0.891 | 0.936 | 0.923 |
| 1D5D | 0.912 | 0.927 | 0.949 |
| 4O37 | 0.190 | 0.215 | 0.936 |
| 1FEV | 0.913 | 0.959 | 0.959 |
| 1D5H | 0.233 | 0.906 | 0.935 |
| 3KMZ | 0.524 | 0.748 | 0.894 |
| 4OKF | 0.843 | 0.948 | 0.935 |
| 1Z3P | 0.870 | 0.945 | 0.951 |
| 1Z3L | 0.120 | 0.940 | 0.936 |
| 4YJW | 0.915 | 0.863 | 0.902 |
| 4LKA | 0.101 | 0.820 | 0.882 |
| 3OQY | 0.665 | 0.845 | 0.847 |
| 1Z3M | 0.122 | 0.934 | 0.942 |
| 5OTT | 0.662 | 0.421 | 0.582 |
| 5OTU | 0.722 | 0.364 | 0.763 |

**Table S9.** DockQ results of linear peptide complexes.

| **PDB ID** | **RFAA** | **NCPepFold w/o Attention** | **NCPepFold** |
| --- | --- | --- | --- |
| 1D5E | 0.825 | 0.850 | 0.850 |
| 6GB1 | 0.946 | 0.838 | 0.892 |
| 3OQZ | 0.795 | 0.864 | 0.841 |
| 3OR0 | 0.814 | 0.860 | 0.907 |
| 2FX8 | 0.000 | 0.000 | 0.081 |
| 2K7L | 0.543 | 0.600 | 0.686 |
| 2RLN | 0.860 | 0.860 | 0.907 |
| 1RBD | 0.884 | 0.860 | 0.884 |
| 1D5D | 0.932 | 0.864 | 0.886 |
| 4O37 | 0.222 | 0.250 | 0.944 |
| 1FEV | 0.936 | 0.936 | 0.957 |
| 1D5H | 0.250 | 0.818 | 0.864 |
| 3KMZ | 0.627 | 0.804 | 0.882 |
| 4OKF | 0.818 | 0.886 | 0.864 |
| 1Z3P | 0.837 | 0.884 | 0.884 |
| 1Z3L | 0.056 | 0.861 | 0.861 |
| 4YJW | 0.894 | 0.851 | 0.872 |
| 4LKA | 0.040 | 0.700 | 0.800 |
| 3OQY | 0.444 | 0.867 | 0.822 |
| 1Z3M | 0.054 | 0.838 | 0.865 |
| 5OTT | 0.829 | 0.683 | 0.659 |
| 5OTU | 0.941 | 0.735 | 0.941 |

**Table S10.** F_nat_ results of linear peptide complexes.

**Section S4. The importance of cyclic positioning**

Figure S1 shows the prediction results visualized with and without the addition of the head-to-tail cyclization matrix. The terminal residues are represented in stick mode. The white sections represent the ground truth structure, the blue sections represent the RFAA predicted structure, and the purple sections represent the NCPepFold predicted structure. RFAA predicted the structure of 2M2X as a linear peptide, whereas NCPepFold successfully incorporated the cyclization information and predicted the head-to-tail cyclic structure. Similar cases are observed for 1T9E, 2J15, 4M6E, and 2M2S. Another example is shown for 2KNN. Although the structure predicted by RFAA looks like a head-to-tail cyclic peptide, the amino and carboxyl groups of the terminal amino acids do not form a peptide bond. Similar cases are observed for 2M1P, 2N8E, 2MSQ, and 2M2G. However, after adding the cyclization constraint, NCPepFold successfully predicts the head-to-tail cyclic peptide structure. The cyclization matrix provides a soft constraint, greatly enhancing the likelihood of cyclization.

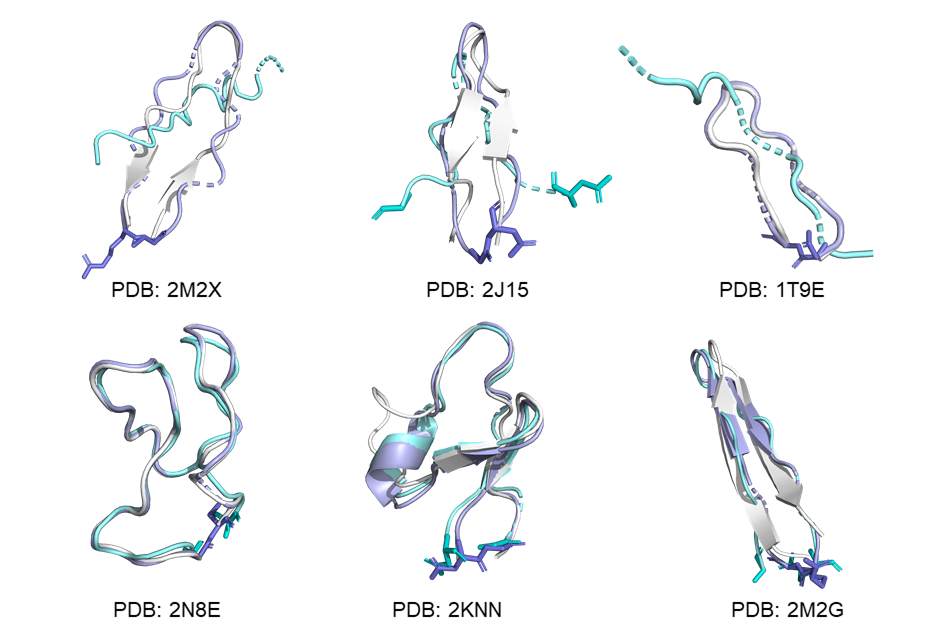

**Figure S1.** Examples of head-to-tail cyclic peptide monomer. The terminal residues are represented in stick mode.

**Section S5.** **Difficult targets**

To evaluate the generalization ability of NCPepFold, we focus on targets with sequence similarity < 25%, referred to as difficult targets. For linear peptide monomers, the difficult targets have the following PDB IDs: 2L87, 2MZA, 2N0N, 2LDA, 3ZS2, 5KGY, 2LDD, 1GEA, 2LDC, 6QXB, 2MYL, 2MZ2, 2MYM. For linear peptide complexes, the difficult targets have the following PDB IDs: 2FX8, 5OTT. For head-to-tail cyclic peptide monomers, the difficult targets have the following PDB IDs: 2N8E, 4M6E, 2J15. For disulfide-bonded cyclic monomers, the difficult targets have the following PDB IDs: 1P9G, 2CRD, 1BIG, 1K64, 2EW4. Table X presents the peptide RMSD and modification RMSD results for each type of peptide, demonstrating that NCPepFold exhibits strong generalization performance.

| **PDB ID** | **Peptide RMSD** | **Modification RMSD** |
| --- | --- | --- |
| **Linear monomer** | | |
| 2L87 | 1.940 | 4.455 |
| 2MZA | 3.156 | 2.139 |
| 2N0N | 1.143 | 3.662 |
| 2LDA | 2.137 | 2.061 |
| 3ZS2 | 3.250 | 2.458 |
| 5KGY | 1.408 | 3.960 |
| 2LDD | 1.364 | 2.380 |
| 1GEA | 3.250 | 1.292 |
| 2LDC | 1.260 | 3.493 |
| 6QXB | 1.115 | 1.956 |
| 2MYL | 1.823 | 2.243 |
| 2MZ2 | 3.323 | 3.451 |
| 2MYM | 2.878 | 2.321 |
| **Head-to-tail cyclic peptide** | | |
| 2N8E | 0.274 | 1.209 |
| 4M6E | 4.063 | 2.090 |
| 2J15 | 1.450 | 0.845 |
| **Disulfide-cyclized peptide monomer** | | |
| 1P9G | 2.104 | 2.301 |
| 2CRD | 1.203 | 1.313 |
| 1BIG | 1.252 | 1.490 |
| 1K64 | 1.740 | 1.132 |
| 2EW4 | 3.587 | 1.558 |
| **Linear complex** | | |
| 5OTT | 5.789 | 4.947 |
| 2FX8 | 12.293 | 1.422 |

**Table S11.** Two RMSD results for peptides with sequence similarity < 25%.
